# Generation of Somatic Mitochondrial DNA-Replaced Cells for Mitochondrial Dysfunction Treatment

**DOI:** 10.1101/2020.10.01.321851

**Authors:** Hideki Maeda, Daisuke Kami, Ryotaro Maeda, Akira Shikuma, Satoshi Gojo

## Abstract

Mitochondrial diseases currently have no cure regardless of whether the cause is a nuclear or mitochondrial genome mutation. Mitochondrial dysfunction notably affects a wide range of disorders in aged individuals, including neurodegenerative diseases, cancers, and even senescence. Here, we present a procedure to generate mitochondrial DNA-replaced somatic cells with a combination of a temporal reduction in endogenous mitochondrial DNA and coincubation with exogeneous isolated mitochondria. Heteroplasmy in mitochondrial disease patient-derived fibroblasts in which the mutant genotype was dominant over the wild-type genotype was reversed over the long term, even inducing the production of pluripotent stem cells from the mitochondrial DNA-replaced cells to maintain the genotype without a reversion to the original. Both mitochondrial disease patient-derived and aged fibroblasts could regain respiratory function and showed lifespan extension. Mitochondrial membranous components were utilized as a vehicle to deliver the genetic materials into endogenous mitochondria-like horizontal genetic transfer in prokaryotes. The mitochondrial DNA-replaced cells could be a resource for transplantation to treat not only mitochondrial diseases, but also senescence-related diseases.

## Introduction

Mammalian mitochondria are composed of more than 1,500 proteins and contain a circular genome, mitochondrial DNA (mtDNA), with several copy numbers (CNs) within a matrix encircled by double membranes and the intermembrane space; mtDNA has been reduced to 16.6 kb during evolution through gene transfer to the nucleus(Nunnari & Suomalainen, 2012). Only 13 mitochondrial proteins are encoded by mtDNA, in addition to 22 tRNAs and 2 rRNAs. Mitochondrial diseases are one of the relatively rare types of inherited metabolic and inflammatory disorders and include two discrete groups that are caused by mutations in mitochondrial or nuclear DNA(Suomalainen & Battersby, 2018). Based on symptoms, not on the types of mutations, mitochondrial diseases are subtyped into a number of syndromes, such as mitochondrial myopathy, encephalomyopathy, lactic acidosis, stroke-like symptoms (MELAS), myoclonic epilepsy with ragged red fibers (MERRF), and Leigh syndrome(Chinnery, 2015). Because mitochondria perform essential functions in cellular physiology, such as energy generation(Smeitink *et al*, 2006), innate immunity(Mills *et al*, 2017), Ca kinetics(Bertero & Maack, 2018), and apoptosis(Burke, 2017), mitochondrial dysregulation and defects lead to various clinical manifestations in many organs at any age, depending on the causative mutations, which may emerge de novo sporadically or are inherited from an autosome, the X chromosome, or maternally.

Mitochondrial diseases with pathological mutations in mtDNA display a phenotype when the rate of mutated versus healthy mtDNA, termed heteroplasmy(Attardi *et al*, 1995; Smith & Lightowlers, 2011), surpasses a given critical threshold (typically 60 – 80%) to biochemical disturbances and defective respiration(Rossignol *et al*, 1999). As the clinical phenotypes of MELAS and MRRF correlate to the heteroplasmy level in muscle, but not in blood(Chinnery *et al*, 1997), there is a heterogeneity of heteroplasmy level between cells even in the same tissue or organ, from organ to organ in the same person(Stewart & Chinnery, 2015). The heterogeneity of mtDNA is divided into intracellular and intercellular heteroplasmy(Aryaman *et al*, 2018), and a single cell analysis in mitochondrial biology has just emerged for clinical samples(Maeda *et al*, 2020b; Walker *et al*, 2020). Current knowledge on mitochondrial diseases has not resulted in the precise prediction and prognosis of patients. Although there has been no curative treatment for mitochondrial diseases and only symptomatic therapy has been provided to patients, understanding of mechanistic insights into the diseases has led to the development of clinical trials using small molecules as well as cell or gene therapy(Russell *et al*, 2020).

At the beginning of the emergence of eukaryotes, an α-proteobacterium was engulfed by the progenitor, enabling evolution(Suomalainen & Battersby, 2018). The capability to take environmental materials up into the cytosol, such as through pinocytosis and especially macropinocytosis, when engulfing large materials is preserved with the current cells and organisms(Amyere *et al*, 2002; Mercer & Helenius, 2009). Horizontal transfer of genetic materials is a common characteristic of bacteria such as *E. coli*; however, these phenomena are also observed in mammalian cells(Pace *et al*, 2008). Since mammalian horizontal transfer of mitochondria through a tunneling nanotube was reported *in an in vitro* experiment (Rustom *et al*, 2004), mitochondrial transfer has been demonstrated with various mechanisms, such as nanotube(Islam *et al*, 2012), extracellular vesicles(Hayakawa *et al*, 2016), and macropinocytosis(Kitani *et al*, 2014), as well as by using an artificial device(Wu *et al*, 2016) and a compound(Maeda *et al*, 2020a). Although the efficiency of mitochondrial transfer did not dominate over the pre-existing mtDNA but provided an additive functionality in the host cells, mechanistic insights of mitochondrial transfer with respect to cell entry and endosomal escape have been deepened(Kami & Gojo, 2020).

The concept of mitochondrial replacement was reported almost three decades ago; this approach utilizes chemicals toxic to mtDNA and enucleated cells as donors of cytosolic organelles, including mitochondria(King & Attardi, 1989). Long-term exposure of cells to ethidium bromide under culture conditions gives rise to mtDNA-free cells, termed ρ0. The *in vitro* enucleation technique(Bayona-Bafaluy *et al*, 2003) enabled the generation of cells with replaced mitochondria, named cybrid cells, through the fusion of ρ0 cells and enucleated cells. *In vivo* mitochondrial replacement in model animals was achieved using either embryonic stem (ES) cell cybrids pretreated with rhodamine 6G or mitochondrial injection into oocytes or zygotes(Nakada & Hayashi, 2011). Gene transfer of endonuclease with a mitochondrial transfer signal to cut and degrade mtDNA offered another ρ0 creation method(Kukat *et al*, 2008). The mitochondria-targeted endonuclease approach has been utilized to generate a heteroplasmy shift of mitochondrial DNA (mtDNA) in mitochondrial diseases(Tanaka *et al*, 2002), rather than to form a deletion of mtDNA for cybrid cells, based on the threshold theory of mitochondrial heteroplasmy(Rossignol *et al*., 1999). Emerging genome editing technology has been extended to the mitochondrial genome to fix a mutation by using mitochondrially targeted Cys2-His2 zinc-finger nucleases (ZFNs)(Gammage *et al*, 2016) and transcription activator-like effector nucleases (TALENs)(Bacman *et al*, 2013). Clinically, mitochondrial replacement therapy (MRT) in assisted reproductive technologies, which is technically executed by pronuclear transfer, maternal spindle transfer, or polar body transfer, is enthusiastic, especially with respect to ethics(Greenfield *et al*, 2017; National Academies of Sciences & Medicine, 2016). Interventions to modulate mtDNA are rising to the level of mitochondrial gene therapy.

Aging alters many biological processes and leads to degenerative disorders, such as Alzheimer’s disease, atherosclerosis, osteoporosis, type 2 diabetes mellitus, and tissue fibrosis, which cause chronic kidney disease and chronic obstructive pulmonary disease(Lopez-Otin *et al*, 2013). A theory that mitochondria drive senescence via reactive oxygen species (ROS), which mitochondria mainly produce, was reported to markedly impact gerontology(Harman, 1956). The theory has been denied in itself; however, the idea that mitochondria play a critical role in cellular senescence, reduced stem cell function and “inflammaging” is currently well accepted(Kauppila *et al*, 2017a). Mitochondria accumulate increased somatic mutations and deletions of mtDNA in various human organs with aging(Corral-Debrinski *et al*, 1992; Cortopassi & Arnheim, 1990), which are directly linked to mitochondrial dysfunction(Fayet *et al*, 2002). Mice defective in a proofreading catalytic subunit of the mtDNA polymerase, called mutator mice, present a short lifespan with premature aging under mtDNA mutation loads(Trifunovic *et al*, 2004). The mutator mouse also displays substantial mitochondrial dysfunction without altering mtROS production(Trifunovic *et al*, 2005). Another mouse is defective in the mtDNA helicase Twinkle exhibited multiple mtDNA deletions, called the deletor mouse, with progressive respiratory dysfunction with time but does not experience accelerated aging(Tyynismaa *et al*, 2005). Conplasmic mice, which possess a C57B/6 nuclear genome and NZB/OlaHsd mtDNA that is different from C57B/6 at 90 positions, exhibit modulated aging processes, consequently resulting in a significant lifespan extension(Latorre-Pellicer *et al*, 2016). Patients suffering from human immunodeficiency virus (HIV) infection and treated with antiretroviral therapy, which inhibits mitochondrial polymerase-γ activity(Sohl *et al*, 2015), demonstrated accelerated aging symptoms, such as increased frailty and augmented cardiovascular diseases(Van Epps & Kalayjian, 2017). Similar to the mutator mouse, treatment with nucleoside analogs for HIV infection provoked mtDNA mutations, the extent of which was proportional to the prescription duration(Payne *et al*, 2011). Although currently, it remains to be elucidated whether the mutations are causative or correlative to aging(Kauppila *et al*, 2017b), a profound mtDNA mutation, at least, can accelerate aging(Jang *et al*, 2018). Replacement of mtDNA in somatic cells could be an intervention to modify aging for mutation overloads in mtDNA.

Since the majority of senescence pathways involve mitochondria, somatic mitochondrial DNA-replaced cells (MirCs) in this study were demonstrated to break the progressively deleterious cycle of senescence and even rejuvenate cells, in addition to the potential to treat mitochondrial diseases.

## Results

### Mitochondrial DNA replacement in normal human dermal fibroblasts

To significantly reduce the mitochondrial genome, a plasmid carrying XbaIR, an endonuclease with a mitochondrial transfer signal, was designed (Figure EV1A). A standard sequence of human mtDNA, referred to as the Cambridge reference sequence (CRS), has five *Xba*IR endonuclease recognition sites (Figure EV1B). The sequence was verified by using isolated human mtDNA, which was digested with *Xba*IR in vitro (Figure EV1C). Mitochondria have a poor capability to repair DNA breaks, and nucleic acid fragments are degraded in the mitochondrial matrix(Tanaka *et al*., 2002). Among the mitochondrial transfer signals that have been reported, we selected the COX8A signal sequence in this study (Figure EV1D), and the ability to label its fusion protein with EGFP in mitochondria was verified in normal human dermal fibroblasts (NHDFs), which were transfected with an electroporator and stained with tetramethylrhodamine methyl ester (TMRM), which is a fluorescent dye that accumulates in mitochondria (Figure EV1E, Fig 1A). The exogeneous endonuclease *Xba*IR, when transferred into mitochondria, effectively reduced the content of mitochondria in immortalized human endothelium EPC100, which is genetically labeled with DsRed2 directed to mitochondria by using the COX8A signal sequence, eliminating the need to use ethidium bromide exposure for ρ0 generation (Figure EV1F, EV1G, and Fig 1B). The mtDNA CN declined to approximately one-third of the original content following gene transfer on day 6, and this cellular state was termed ρ(-) cells (Fig 1C). The mRNA expression analysis of *Xba*IR showed time-dependent expression with a peak on day 2 and a rapid decrease to undetectable levels on day 14 (Figure EV1H). Puromycin exposure for 24 h could enrich the gene-transferred cells (Figure EV1I, and EV1J). On day 12 following puromycin selection, the mitochondrial membrane potential, a surrogate marker of mitochondrial genome reduction, showed a clear decrease in NHDFs (Figure EV1K). The protocol to replace the mitochondrial genome, which constitutes a 1-day drug selection on day 2 and cocultivation with isolated exogenous mitochondria on day 6, was designed (Figure EV2A). On day 12 following coincubation with isolated mitochondria, the content of mitochondria recovered to the original level (Fig 1D). Recently, the simple centrifugation of a mixture of cells and isolated mitochondria was reported to increase the mitochondrial uptake of cell lines, especially cancer cells, just 2 days after the procedure(Kim *et al*, 2018). However, we found that only a small increase in uptake into primary fibroblasts occurred with the procedure (Fig 1E). This may be attributed to the decreased macropinocytosis in primary cells compared to cancer cell lines. NHDFs treated with the mitochondria replacement protocol were examined by sequencing short mitochondrial genome fragments with the difference between the recipient, NHDF, and donor, EPC100 (Figure EV2B). Both nucleotides originating from EPC100 and NHDF were significantly present on day 12 after mitochondrial replacement, although PCR-directed sequencing could not be quantitatively analyzed (Figure EV2C). A TaqMan single nucleotide polymorphism (SNP) genotyping assay was applied to quantify the origin of the mitochondrial genome (heteroplasmy) based upon the difference at position 16362 (NHDF: T, EPC100: C) in the D-loop (Figure EV2D). The generated cells, named <u>MirCs</u>, contained mitochondrial genomes with more exogeneous genes than endogenous genes (Fig 1F), suggesting that this protocol could revert the heteroplasmy of cells in mitochondrial diseases.

**Figure 1:**
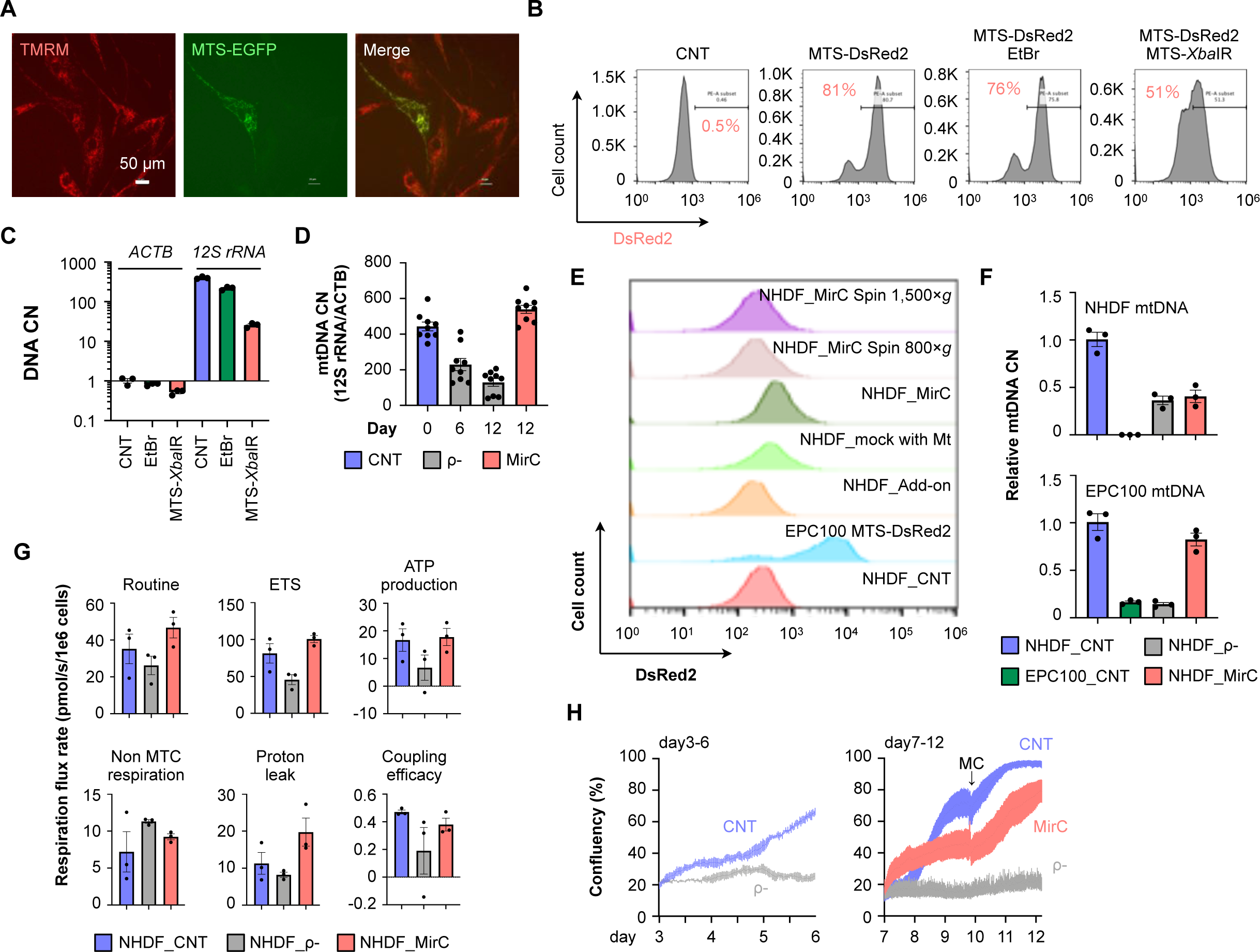
MirCs were generated from normal human dermal fibroblasts (NHDFs). (A) Left; TMRM staining specific for mitochondria in NHDFs, middle; NHDFs, which were transfected with pCAGGS-MTS-EGFP-PuroR and selected with puromycin, reticularly expressed, right; EGFP completely merged the red fluorescence of TMRM, verifying that MTS correctly transported downstream genes of interest. White bar indicates 50 μm. (B) The reductions in DsRed2 in endonuclease-treated cells were analyzed with FACS. (C) By qPCR on day 5 following the treatment for mtDNA reduction, the mtDNA of endonuclease-treated cells was decreased to approximately 60%, clearly contrasted with the effect of EtBr treatment. (D) Mitochondrial copy number (mtCN) following mtDNA replacement, showing a clear increase in mtCN to baseline levels on day 12 with coincubation of isolated mitochondria. (E) Comparison of mitochondrial transfer with other procedures that use centrifugation. DsRed2-labeled mitochondria were used as donors and analyzed on day 2 by FACS. Only endonuclease-treated cells engulfed a significant number of exogeneous mitochondria. (F) Heteroplasmy in MirC-derived NHDFs and mitochondria of EPC100 cells showed the domination of exogeneous origin with a minor portion of the endogenous genotype. (G) Key parameters of respirometry were quantified. (H) Although the proliferation of MirCs was lower than that of the parental cells from day 6 to day 9, that of MirCs was equivalent to that of the parental cells after day 9. ρ(-) cells quite slowly proliferated throughout all periods. MC: medium change.

### Metabolic recovery of MirC from ρ(-) cells

Whether the transferred mtDNA could generate energy was investigated with a coupling control protocol by using high-resolution respirometry. Representative oxygen consumption rate curves of native cells, ρ(-) cells, and MirCs on day 12 are shown, and the respiratory flow and control ratio were calculated (Figure EV2E, Fig 1G). Routine respiration, electron transfer system (ETS) maximum capacity, ATP production, and coupling efficiency showed the same trends: the indexes were decreased in ρ(-) cells compared with parental cells and increased in MirCs, approaching the levels in parental cells. Proton leakage was increased following mtDNA replacement compared with the native and ρ(-) cells, which might reflect mitochondrial membrane damage under recovered respiration. In addition, the phenotypic recovery of MirCs was demonstrated with respect to cell proliferation, while ρ(-) cells showed a poor proliferative capability (Fig 1H). These results confirmed that this protocol can enable the replacement of mtDNA in somatic cells with clinically applicable materials and procedures.

### Mitochondrial DNA replacement in fibroblasts in a mitochondrial disease patient, Leigh syndrome

We next attempted to correct cells derived from patients with mitochondrial disease with mtDNA mutations by using the somatic mtDNA replacement protocol. We used primary fibroblasts derived from a patient diagnosed with Leigh syndrome with a C10158T mutation when she was 3 months old, named 7S fibroblasts (Figure EV3A, EV3B). The same mtDNA replacement protocol performed with NHDFs was applied to 7S fibroblasts. The sequencing of mtDNA at the 10158th nucleotide showed C in the donor mitochondria derived from EPC100 as CRS (Figure EV3A). The mtDNA CN assay showed that the kinetics following the mtDNA replacement protocol were almost the same as the kinetics observed in NHDFs (Fig 2A). Differences were observed in the reduction rate, which was decreased significantly more in 7S fibroblasts than in NHDFs both 6 days and 12 days after the mtDNA replacement protocol. The mtDNA content recovered to the baseline level in the original 7S fibroblasts on day 12. Whether the mitochondria in 7S fibroblasts contain exogeneous mtDNA was examined by sequencing the mitochondrial genome fragment including the 10158th nucleotide. Large waves of C and small waves of T were observed in the cells that received mitochondrial replacement (Figure EV3C). We performed a TaqMan SNP genotyping assay to estimate heteroplasmy through this intervention (Figure EV3D). The original heteroplasmy in 7S fibroblasts was more than 90%, whereas the MirCs derived from 7S fibroblasts exhibited approximately 20% heteroplasmy on day 12 after mitochondrial transfer (Fig 2B, Figure EV3E). We investigated heteroplasmy in single cells by using single-cell droplet digital PCR, which enabled multiplex detection for the presence or absence of a particular genotype in a cell (Fig 2C). The results revealed that the level of intracellular heteroplasmy changes at the single-cell level. The results revealed that the level of intracellular heteroplasmy changes at the single-cell level. MirCs derived from 7S fibroblasts regained the proliferation capability in ρ(-) cells, although the proliferation rate of MirCs was still less than that of the parental 7S fibroblasts (Fig 2D). For long-term culture, MirCs demonstrated dramatic lifespan extension up to the 63rd population doubling level (PDL), given that growth arrest occurred when the doubling time was more than 120 h (Fig 2E, 2F). The cells received mitochondrial replacement at the 8th PDL so that the reconstituted cells with healthy mtDNA might have a lifespan up to the 55th PDL, which is the Hayflick limit in cell culture. A mutation in the coding sequence of the ND4 gene of Complex I in the respiratory chain in 7S fibroblasts impaired the ability of Complex I to transfer electrons coupled with pumping protons up from the matrices to the intermembrane space. Glycolysis was the dominant mitochondrial ATP production pathway in 7S fibroblasts, resulting in the compensatory adaptation of a larger cell size to contain more mitochondria even with damage and poor efficacy (Fig 2G). The diameter of 7S fibroblasts was approximately 1.5 times larger than that of NHDFs by the 10th PDL and increased by an additional 2 times by the 20th PDL, so the volume was 3 to 8 times larger. In line with the proliferation recovery after mitochondrial replacement with 7S fibroblasts, the cellular size decreased over time to that of NHDFs at a comparable PDL. The incorporation of the donor cell line EPC100 into MirCs was decreased by using a short tandem repeat (STR) assay (Fig 2H), and the tumorigenesis of MirCs was denied by measuring the expression of TERT as a tumor marker (Fig 2I).

**Figure 2:**
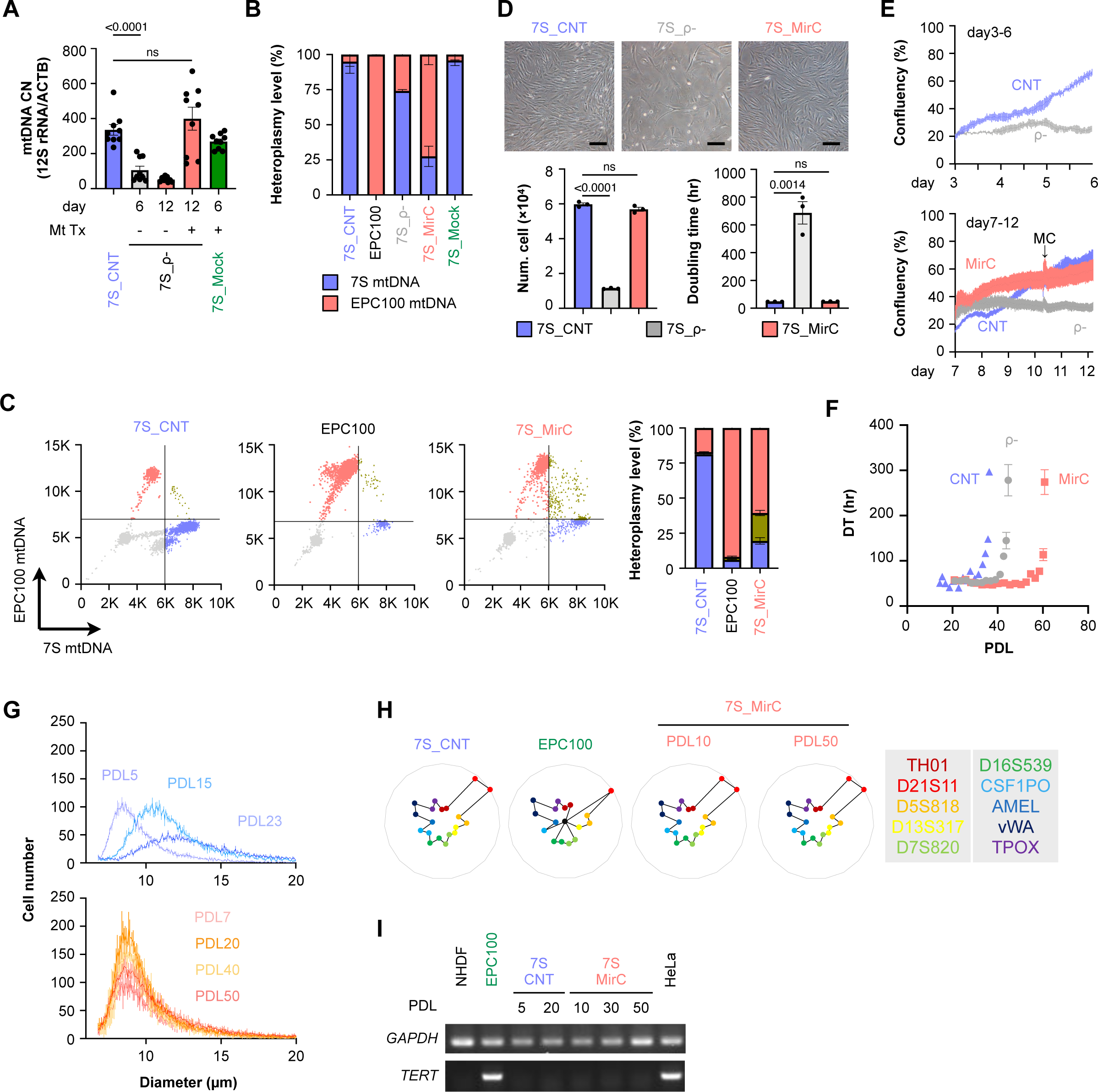
MirCs were generated from mitochondrial disease patient-derived (7S) fibroblasts. (A) During the procedure, mtDNA CN. (B) TaqMan qPCR SNP genotyping assay demonstrated the dominance of exogenous mtDNA. (C) Single-cell ddPCR analysis for heteroplasmic MirCs. (D) Microscopic photographs of cell cultures following mitochondrial replacement. The black bar indicates 200 μm. (E) Cell growth of MirCs compared with the original cells and ρ(-). (F) Long-term culture showed the lifespan extension of MirCs. (G) The cell size of MirCs was maintained, whereas that of the original cells was significantly enlarged from early PDL. (H) Short tandem repeats demonstrated no contamination of the original MirCs. (I) TERT expression in MirCs. mtDNA, mitochondrial DNA. CNT, no treatment control cell. ρ-, rho minus, indicates cells with a low mtDNA number. CN, copy number. MC, medium change. DT, doubling time. PDL, population doubling level.

### Phenotypic recovery in MirC derived from fibroblasts of a mitochondrial disease patient

The concentration of citrate synthetase (CS), which is used to standardize cellular content, was 2 times higher in 7S fibroblasts by the 10th PDL than in NHDFs. MirCs derived from 7S fibroblasts were investigated by using Oroboros O2k with respect to respiratory function over time after the transfer (Fig 3A). Routine respiration and free routine activity (ATP production) decreased by the 20th PDL after replacement, and the maximum capacity of the ETS maintained the original levels of 7S fibroblasts. On the 30th PDL after the replacement, all three indexes with respiratory function strikingly ascended and surpassed the original (Fig 3B, 3C). These results indicate that it takes a given time period to reconstitute the electron transfer system with healthy and nonmutated Complex I following mitochondrial replacement. In the early phase after the replacement, a reduction in mutated mtDNA might have exposed a more critical energy shortage than that in NHDFs; consequently, the delay in recovery might have emerged. Residual oxygen consumption (ROX) decreased over time after replacement. The reason why the change occurred earlier than that of ATP production might be related to the decrease in cell size (Fig 2G), which could have reduced the energy demand. Leak respiration, which reflects intrinsic uncoupling, such as proton leakage, proton slip, and electron slip, decreased during a shortage of ATP production in mitochondria and maintained low values at the time when the reconstitution was likely complete. The kinetics of the coupling rate were the same as the kinetics of ATP production, consistent with the strong relationship between the coupling rate and ATP production. The functional reversal on the 30th PDL after replacement, which was comparable with that of NHDFs, indicates that the strategy could have potential clinical applications. Flux control ratios (Fig 3C) independent of cell size and mitochondrial content were calculated. ROX/E’ indicates the shift to mitochondria with respect to oxygen consumption, L/E indicates the amelioration of the decoupling of electron transfer systems, R/E indicates the improvement in respiratory capacity by reconstituting Complex I with the wild-type complex, and (R-L)/E represents phosphorylation-related respiration, indicating metabolic recovery at the late stage. The stress responses of 7S fibroblasts were examined with an oxidative stress model exposed to H_2_O_2_ and a starvation model cultivated in essential amino acid-free medium to evaluate apoptosis. By using Annexin V as an early apoptosis marker for phosphatidyl serine and propidium iodide as a late apoptosis or necrosis marker, the levels of both stresses, in which NHDFs were tolerated and 7S fibroblasts were significantly damaged, resulting in apoptosis, were increased (Fig 3D). MirCs derived from 7S fibroblasts were protected from both stresses with similar apoptosis levels to NHDFs. Although IL-6 increased among senescence-associated secretory phenotypes in 7S fibroblasts, MirCs derived from 7S fibroblasts showed restored IL-6 expression (Fig 3E). These results might suggest that sensitivity to cell death and progeric phenotype in 7S fibroblasts could also be reverted to normal through mtDNA replacement.

**Figure 3:**
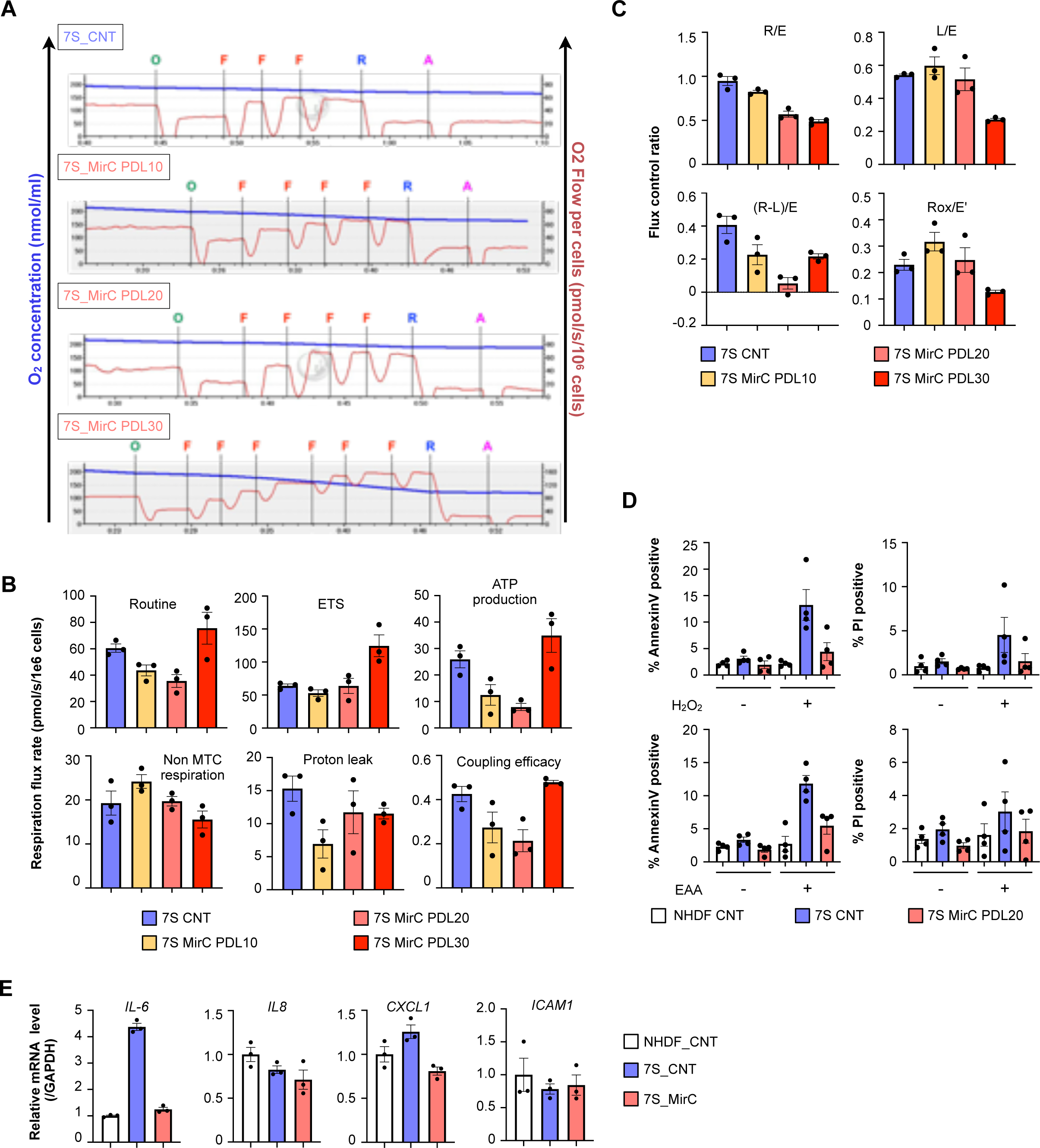
Phenotypic characterization of MirCs. (A) Respiratory functions of MirCs at PDLs 10, 20, and 30 following mtDNA replacement by using a coupling control protocol in Oroboros O2k, O, 4 mg/ml oligomycin. F, 1 mM carbonyl cyanide 4-(trifluoromethoxy)phenylhydrazone (FCCP). R, 1 mM rotenone. A, 5 mM antimycin. (B) Respiratory flow rates demonstrated that it took approximately 20 PDLs to improve respiratory function. (C) Flux control ratios were ameliorated earlier than respiratory flow rates. (D) Mitochondrial disease patient-derived fibroblasts were more sensitive to H_2_O_2_ and essential amino acid (EAA)-free medium than their MirCs. (E) Compared with the parental cells, MirCs showed significantly suppressed IL-6, representative factors of the senescence-associated secretory phenotype. CNT, no treatment control cell. PDL, population doubling level. ETS, electron transport system. Non-MTC respiration, nonmitochondrial respiration.

### Kissing exogeneous mitochondria with endogenous mitochondria to transfer donor mtDNA

The fate of the mitochondria transferred into cells was investigated separately on the outer membrane, inner membrane, or nucleoid, including mtDNA. Mitochondrial transcription factor A (TFAM), which binds to mtDNA and regulates mitochondrial biogenesis, was selected to trace exogeneous mitochondrial nucleoids. Genetically, TFAM-marked cells were created by infecting the recombinant retrovirus carrying the sequence coding a fusion protein with TFAM and EGFP (Figure EV4A to EV4C). TFAM-marked mitochondria were used as donors, and DsRed2-marked cells in the mitochondrial matrices were used as recipients. During the mtDNA replacement protocol, cells were observed by hyperfine microscopy. The donor nucleoids settled in the pre-existing mitochondrial matrices (Fig 4A). In the combination of DsRed2-marked mitochondria derived from EPC100 as donors and ρ(-) cells derived from genetically EGFP-marked NHDFs as hosts, the fate of mitochondrial protein components was examined using superfine microscopy (Fig 4B). The contacts of the donor and the resident mitochondria were recognized, but no broad fusion was observed (Fig 4B, Movie EVs 1-1 to 1-4). Moreover, the fluorescent spots of donor mitochondria on day 2 following isolated mitochondrial coincubation were quite rare, suggesting that exogeneous mitochondrial protein components might be degraded for a few days.

**Figure 4:**
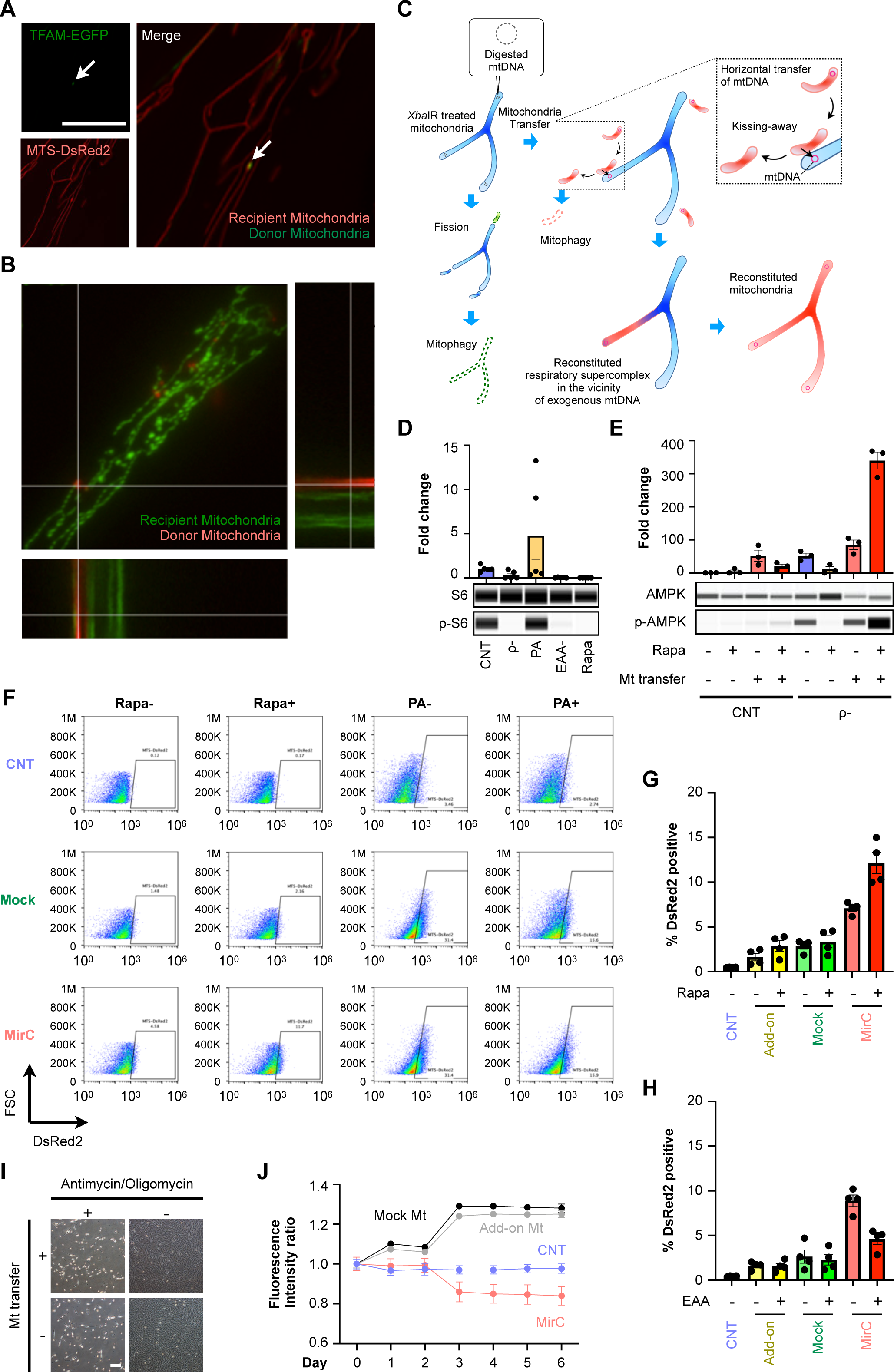
Mechanistic insights into the process of mtDNA replacement. (A) Superfine microscopy showed that exogenous mtDNA coupled with TFAM existed in endogenous mitochondria. White bar indicates 50 μm. (B) Two fluorescence signals were always far apart and showed temporary contact for a short period. (C) A model of “Kissing-away” in transferring mtDNA into endogenous mitochondria. (D) Activation of mTorc1 was measured by the ratio of phosphorylation of S6 to S6. ρ(-) was significantly decreased, similar to that observed in essential amino acid-free medium and rapamycin treatment. (E) AMPK activation during the MirC generation protocol. (F-H), By measuring fluorescence that marked donor mitochondria in FACS, rapamycin, an mTorc1 inhibitor, enhanced the engulfment of exogenous mitochondria, regardless of preconditioning, and palmitic acid suppressed it only in the MirC generation protocol. (I) ρ(-) with a minimal amount of mtDNA could ingest exogenous isolated mitochondria, whereas ρ0 without mtDNA could not, suggesting that macropinocytosis should be supported by ATP generated by remnant mitochondria. White bar indicates 200 μm. (J) Fluorescent area was measured as a surrogate for the fate of exogeneous mitochondrial membranous components. PA, palmitic acid.

These results lead to a model to transfer mtDNA in this protocol that engulfed exogeneous mitochondria with mtDNA fuses with pre-existing mitochondria that lost mtDNA, mtDNA is transferred to the pre-existing mitochondrion, and the vacant mitochondrion is degraded (Fig 4C).

### Macropinocytosis of exogeneous mitochondria is regulated by the mTORC1 pathway

We investigated the mechanism that regulates the macropinocytosis of exogeneous mitochondria to create an ideal protocol to generate MirCs of any type of cell. Since ρ(-) cells are exhausted of ATP to support cellular activities, the intracellular energetic state of ρ(-) cells could be compatible with starvation. Essential amino acid-free medium for AMPK stimulation, palmitic acid (PA) for mTORC1 activation, and rapamycin for mTORC1 suppression were chosen to examine the state of ρ(-) cells. The ratios of phosphorylated AMPK to AMPK and phosphorylated p70 S6 kinase to p70 S6 kinase, which is a downstream target of mTORC1, were examined by using capillary electrophoresis, namely, Wes™ (Protein Simple) (Fig 4D, 4E). The AMPK pathway was significantly activated in ρ(-) cells, and the mTORC1 pathway was drastically suppressed in ρ(-) cells to the same level as that observed under starvation conditions or rapamycin treatment (Fig 4D, 4E). AMPK activation in ρ(-) was increased with the coincubation of exogeneous mitochondria and was further increased with the addition of rapamycin (Fig 4F). We examined the effects of rapamycin and PA on mitochondrial engulfment during mitochondrial cocultivation. On day 8, FACS analyses were performed to detect DsRed2, the donor mitochondria. Rapamycin significantly augmented the engulfment of the mitochondria, whereas PA clearly suppressed it (Fig 4F to 4H). In both mock transfectant and add-on type mitochondrial transfer, there were no significant differences. This suggests that mTORC1 could be a regulator of mitochondrial macropinocytosis. Rapamycin, a drug used in the clinic, can be utilized to increase the efficiency of macropinocytosis in cells with poor macropinocytosis. Whether the mtDNA replacement protocol can be refined through the complete elimination of endogenous mtDNA(Correia-Melo *et al*, 2016) was examined by using cells genetically modified to overexpress PARKIN, which is a key molecule for mitophagy (Fig 4I). The transfectants can expose ρ0 to oligomycin and actinomycin, and the ρ0 derived from the transfectants could not engulf exogeneous mitochondria and showed poor mobility (Movie EV 2-1 to 2-4). The ρ(-) NHDFs (Movie EVs 3-1 to 3-3) and 7S fibroblasts (Movie EV 4) markedly engulfed the isolated mitochondria, as observed by time-lapse microscopy. However, both naïve NHDFs (Movie EV 2-2) and mock-transfected NHDFs (Movie EV 2-3) seemed to gather the isolated mitochondria, forming very large aggregates on the cell surfaces. The decline in engulfed exogeneous mitochondria started as soon as 72 h following the coincubation in the mtDNA replacement protocol (Movie EV 2-1), whereas the fluorescence of exogeneous mitochondria in either the mock transfectants or mitochondria add-on protocol increased rather than decreased (Fig 4J). In our previous study, electron microscopy showed mitochondria of high quality, but some fragments seemed to be dissociated from mitochondria, and some damaged mitochondrial membranes were observed(Kitani *et al*., 2014). A partial reduction in mtDNA could be essential for this protocol because a complete reduction inhibits macropinocytosis of exogeneous mitochondria, and add-on mitochondrial coincubation does not efficiently execute macropinocytosis.

### No reversion of heteroplasmy in induced pluripotent stem cells derived from MirC

Although there are many reports on the successful generation of induced pluripotent stem (iPS) cells from cells derived from patients with mitochondrial diseases(Hsu *et al*, 2016), we could not generate iPS cells from 7S fibroblasts by using standard methods of Sendai virus(Ban *et al*, 2011) carrying OCT3/4, SOX2, KLF4, and c-MYC despite several attempts (Fig 5A to 5D). We acquired several lines of iPS cells from MirC-derived 7S fibroblasts (Fig 5C to 5G). These iPS cells possess approximately half of the parental cells without any variability (Fig 5G), consistent with reports that embryonic stem cells have a lower mtDNA CN than parental differentiated cells(Lonergan *et al*, 2007). The MirC-derived iPS cells exhibited less than 10% heteroplasmy, which is similar to the heteroplasmy in the parental MirCs on day 160 following gene transfer (Fig 5F, 5G). Mitochondrial gene replacement (MR) in oocytes can provide offspring(Tachibana *et al*, 2009), and MR in pluripotent stem cells with mitochondrial dysfunctions can normalize their metabolism(Ma *et al*, 2015). Although MR therapy has been applied in the clinic(Greenfield *et al*., 2017), concerns that the mitochondrial genotype can be reverted to the original following MR have been raised(Hudson *et al*, 2019). A mechanistic study on reversion revealed that genetic drift could be the cause(Yamada *et al*, 2016). The mtDNA replacement protocol in somatic cells did not show reversion in long-term cultivation and maintained the original heteroplasmy level.

**Figure 5:**
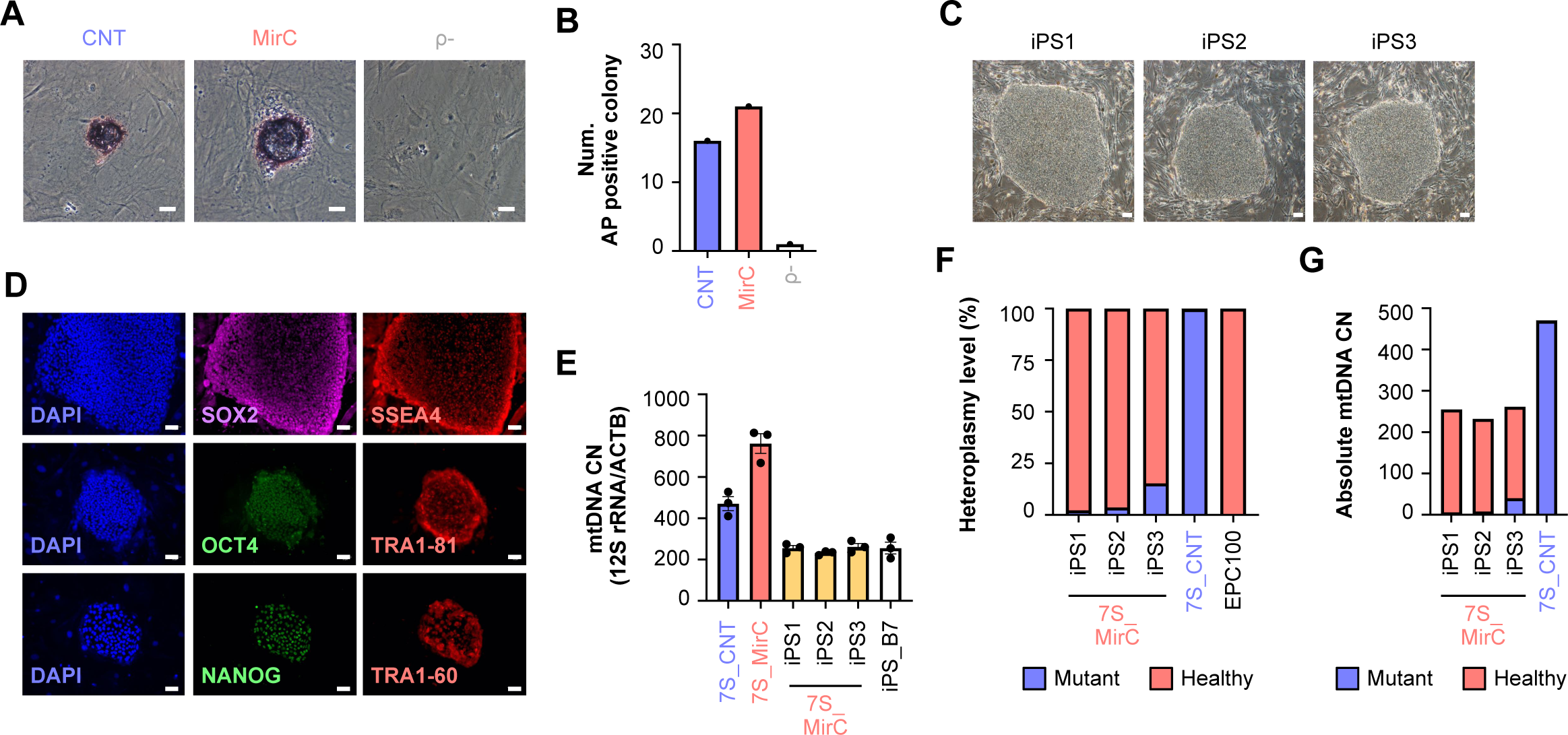
Characterization of induced pluripotent stem cells derived from the MirCs of mitochondrial disease patient fibroblasts. (A), (B) Alkaline phosphatase assay for initial colonies showed poorly formed colonies in diseased fibroblasts and firm colonies in MirCs. White bar indicates 50 μm. (C) Representative photographs of iPS colonies in the long term. White bar indicates 100 μm. (D) Expression of stem cell-related transcription factors, such as SOX2, OCT3/4, NANOG, SSEA4, TRA1-81, and TRA1-60. White bar indicates 50 μm. (E) MtDNA CNs in iPS cells derived from MirCs of mitochondrial diseased fibroblasts. (F), (G) Heteroplasmy in iPS cells maintained the original rate of MirCs without reversion to the diseased parental rate. The replacement of mtDNA affected the lifespan of cultured fibroblasts.

### Lifespan extension of cultivated fibroblasts by mtDNA replacement

To examine the hypothesis that mtDNA replacement can rejuvenate senescent cells, two models were designed to examine the cell cycle, namely, doubling time and PDL. NHDFs with early PDL (approximately 5 to 10, called “young”) or late PDL (approximately 40 to 45, called “aged”) were utilized for the two models. One model involved young cells containing mitochondria derived from old cells, designated “A2Y,” and the other model involved aged cells containing mitochondria derived from young cells, designated “Y2A” (Figure EV5A). The Y2A model clearly demonstrated an increase in lifespan to approximately 65 PDLs (Fig 6A). Control old cells and mock transfectant cells showed growth arrest at 55 PDLs, which is consistent with the Hayflick limit. On the other hand, the A2Y model demonstrated a reduced lifespan of approximately 45 PDLs (Fig 6B). The changes in the cellular lifespans in both models could be attributed to exogenous mtDNA. The endogenous lysosomal β-galactosidase that accumulates in senescent cells was reduced in Y2A cells but not completely restored to the level observed in young cells (Fig 6C, 6D). Respiratory function analysis of MirCs in the Y2A model demonstrated that routine respiration, ETS, spare respiration, and ATP production were significantly increased, and nonmitochondrial respiration was significantly decreased but did not reach the level in young cells (Fig 6E, Figure EV5B to EV5D). Unaffected proton leakage might be attributed to incomplete reconstitution by exogeneous mtDNA on day 6 following mitochondrial coincubation. These results demonstrate that the replacement of mtDNA in old cells with young cell mtDNA could provide a rejuvenation modality.

**Figure 6:**
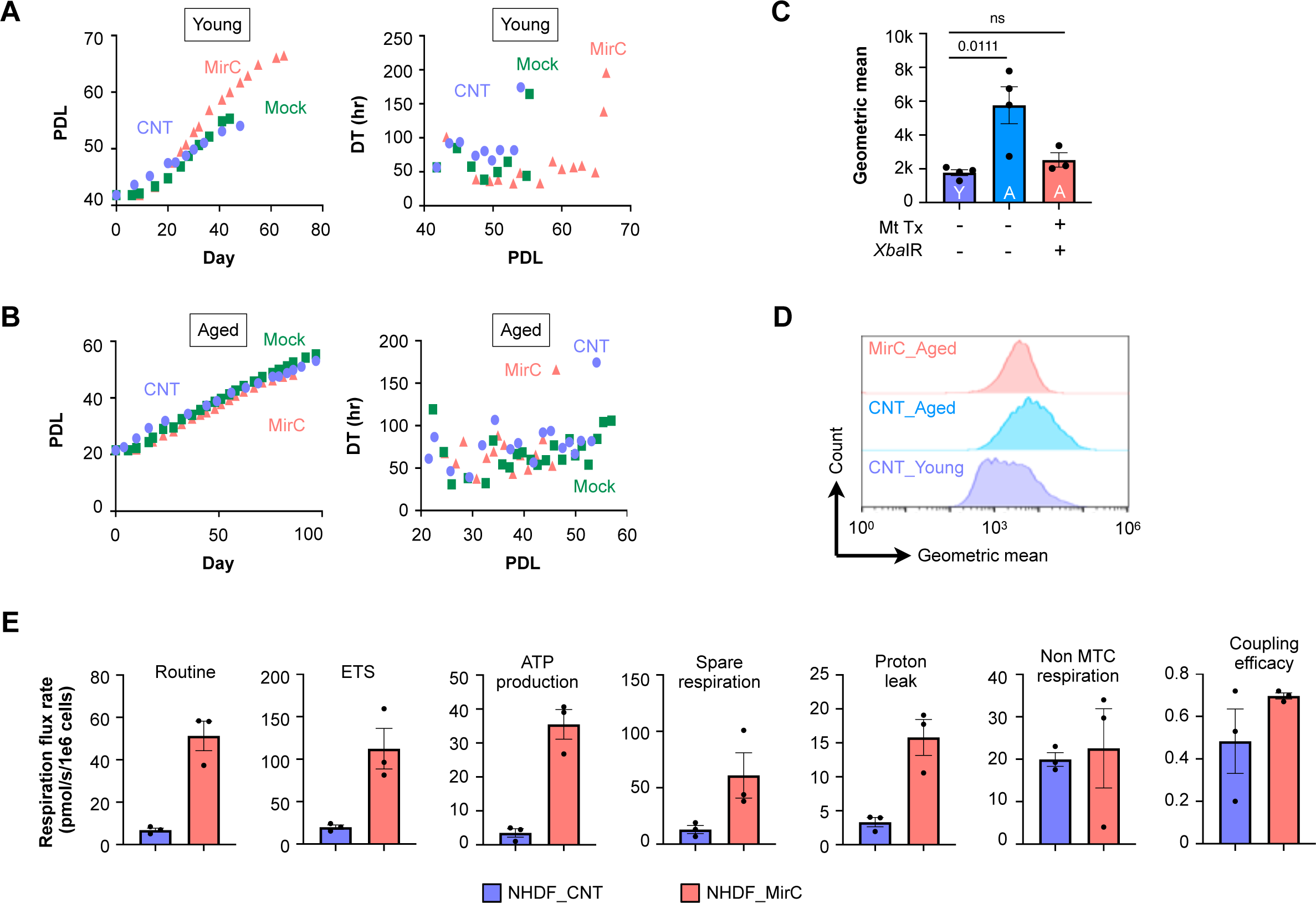
Lifespan extension of cultivated fibroblasts by mtDNA replacement. (A), (B) Normal human dermal fibroblasts with high population doubling levels, approximately 40 (aged NHDFs), that converted MirCs with mitochondria-isolated NHDFs with low PDLs, approximately 10 (young NHDFs), showed an extended lifespan until growth arrest, whereas MirC-derived NHDFs derived from young NHDFs showed a shortened lifespan. (C), (D) β-Gal staining in Y2A MirCs decreased compared with the original and mock transfectants, as evidenced by FACS analysis. (E) Respiratory flow rates of Y2A MirCs significantly improved from the original aged NHDFs.

## Discussion

Isolated mitochondria are mainly engulfed via macropinocytosis(Kitani *et al*., 2014), although other processes are involved in the internalization of exogeneous mitochondria. The induction of macropinocytosis takes place by either growth factors, pathogens including viruses or bacteria, or apoptotic bodies(Mercer & Helenius, 2009). It was reported that macropinocytosis and mTORC1 share signaling pathways and seem to coordinately contribute to cellular growth(Yoshida *et al*, 2018). We focused on two molecular pathways, mTORC1 and AMPK; the former acts as an essential sensor of amino acids, energy, oxygen, and growth factors and a key regulator of protein, lipid, and nucleotide synthesis(Zoncu *et al*, 2011); the latter is a sensor of AMP levels whose activation results in autophagy, mitochondrial biogenesis, glycolysis, and lipolysis(Herzig & Shaw, 2018). Both pathways are involved in the uptake of extracellular nutrients. Macropinosomes that are released into the cytosol following the engulfment of extracellular materials fuse to lysosomes and lead to mTORC1 recruitment and activation(Yoshida *et al*, 2015). Activated mTORC1 negatively regulates macropinocytosis through insulin response substrate (IRS)(Harrington *et al*, 2005). One report showed increased macropinocytosis in Ras-overexpressing cancerous cells upon the addition of rapamycin(Sung *et al*, 2015). On the other hand, under a nutrient-replete state, an inhibitor of mTORC1 was reported to suppress macropinocytosis(Hackstein *et al*, 2002). Moreover, during starvation, mTORC1 suppression stimulates the degradation of proteins in lysosomes to survive without affecting the macropinocytosis machinery(Palm *et al*, 2015). Macropinocytosis is the molecular underpinning of exogenous mitochondrial transfer, and ρ(-) cells, whose mitochondrial function is severely impaired, metabolically deviate to glycolysis in the presence of rich nutrients, which is different from any previous studies, resulted in enhanced macropinocytosis. Upon ρ(-) cells and the presence of isolated mitochondria, rapamycin is an effective activator for macropinocytosis. Pharmacological activation of AMPK in macrophages increased macropinocytosis(Guest *et al*, 2008). AMPK is regulated by not only the tumor suppressor LKB1 in response to mitochondrial insults and energy stress(Shackelford & Shaw, 2009) but also the calcium-sensitive kinase CAMKK2 in response to hormones and stresses that acutely modulate intracellular calcium(Herzig & Shaw, 2018). Zaire Ebolavirus requires AMPK activation upon glycoprotein-mediated macropinocytic entry(Kondratowicz *et al*, 2013). Mitochondrial DNA reduction in the protocol generating MirCs is a potent activator for AMPK with an increased AMP-to-ATP ratio, relayed to enhanced macropinocytosis for exogenous mitochondria.

Pathogens to be engulfed in macropinosomes can evade the innate immune response, as engulfed apoptotic bodies in macropinosomes do not induce innate immunity(Albert, 2004). Although solutions of isolated mitochondria contain some damaged mitochondrial membrane fragments and genomes, called mitochondrial damage-associated molecular patterns, which might induce innate immunity in the cytosol(Grazioli & Pugin, 2018), the engulfed mitochondria in macropinosomes have to circumvent the activation of innate immunity and escape to the cytosol to function in the new destiny. Following the internalization of mitochondria by macropinocytosis, how exogenous mitochondria break macropinosomal membranes has not yet been revealed, although our results indicate that exogenous mitochondria transfer their genome to endogenous mitochondria. It is essential for entrapped pathogens in endosomes to break through the membranes of endosomes to the final destination, and pathogens take advantage of various machineries, such as membrane fusion, pore formation, receptor binding, proteolytic processing, and intraluminal acidification(Staring *et al*, 2018). In addition, mitochondria form microdomains with endosomes to transport iron and cholesterol, which are indispensable for the synthesis of iron-sulfur clusters and steroid hormones, respectively(Charman *et al*, 2010; Sheftel *et al*, 2007). Packaged mitochondria, such as either damaged or healthy mitochondria, enclosed by the membrane, called mitochondria-derived vesicles, attack endosomes containing Staphylococcus aureus by releasing ROS(Abuaita *et al*, 2018) and are exported from the cell to be utilized to rescue growth in respiration-deficient tumors(Dong *et al*, 2017). Either direct contact of macropinosomes holding exogenous mitochondria with endogenous mitochondria or breakage of the macropinosomal membrane might occur prior to the transport of exogenous mitochondrial DNA to the host mitochondria.

Hallmarks of aging have been classified into nine categories, all of which are key players in mitochondria(Lopez-Otin *et al*., 2013). The activated DNA repair machinery upon genomic damage influences mitochondrial homeostasis with raised mitochondrial ROS production, increased membrane potential, and limited mitophagy through inhibition of the NAD^+^-dependent deacetylase sirtuin 1 (SIRT1), resulting in the accumulation of dysfunctional mitochondria(Fang *et al*, 2014). Half a century ago, Hayflick discovered that cultivated cells have a limitation of doubling, which is 50 to 55 PDLs(Hayflick, 1965). One of the molecular bases of the Hayflick limit is the length of telomeres(Harley *et al*, 1990), which are shortened in senescence but are catalyzed and lengthened by telomerase, which bypasses senescence and promotes growth indefinitely(Bodnar *et al*, 1998). Telomere attrition activates p53, which leads to the suppression of Pgc1α/β, nuclear respiratory factor 1, estrogen-related receptor α, and Pparα, consequently resulting in inhibition of mitochondriogenesis and mitochondrial dysfunction(Sahin *et al*, 2011). Epigenetic alterations in aging are affected by acetyl-coenzyme A, which is generated in mitochondrial matrices and utilized as a donor of chromatin acetylation(Su *et al*, 2016). Impaired proteostasis(protein homeostasis) is another characteristic of aging(Labbadia & Morimoto, 2015). The integrity of proteostasis is strictly maintained by translational regulation, protein folding chaperone, ubiquitin-proteosome system (UPS), and the autophagy-lysosome system (ALS)(Vilchez *et al*, 2014). Because chaperones depend upon ATP, a decrease in bioenergy with aging jeopardizes correct protein folding(Brehme *et al*, 2014). Both UPS(Chondrogianni *et al*, 2015) and ALS, including mitophagy(Madeo *et al*, 2015), decline with time. The alternations of these three systems generate aggregates not to be recycled in the cytosol and lead to degenerative disorders. MirCs derived from aged fibroblasts that received mitochondria in young fibroblasts exhibited a significant regain of OXPHOS ability and cumulative proliferation, suggesting that the abovementioned primary aging hallmarks might be reversible with mitochondrial DNA replacement.

Mitochondrial dysfunction might be a main player as not only a consequence but also a causative factor in the senescent phenotype with chronic inflammation(Wang & Hekimi, 2015). In contrast, a mild inhibition of mitochondrial respiration extended in *C. elegans* through mtROS signaling that increased HIF-1 activity(Lee *et al*, 2010) and mitochondrial unfolded protein response(Yee *et al*, 2014). Although this evidence alerts an oversimplification that the more highly active mitochondria, the longer lifespan(Theurey & Pizzo, 2018), malfunctions of mitochondria too harsh to enable compensation response, which are observed in mutator mice and fibroblasts derived from ND5-deficient mitochondrial disease in this experiment, could be attributed to premature aging. Deregulated nutrient sensing, in which a shortage of nicotinamide adenine dinucleotide (NAD^+^) caused by downregulation of nicotinamide phosphoribosyl transferase (NAMPT) and hyperactivation of poly(ADP-ribose) polymerase 1 (PARP1) leads to inhibition of NAD^+^-dependent deacetylase sirtuin 1 (SIRT1), relays to the acetylation-dependent inactivation of PGC1α, consequently resulting in depressed biogenesis of mitochondria that exaggerates NAD^+^ availability(Gomes *et al*, 2013). Low PGC1α activity not only downregulated the expression of mitochondrial proteins encoded in the nucleus but also of the mitochondrial transcription factor TFAM complexing with mtDNA. In addition to two core senescence-regulating pathways, p53 and p16/Rb, the senescence-associated secretory phenotype (SASP) (Coppé *et al*, 2008; Kuilman & Peeper, 2009), in which an array of inflammatory cytokines, chemokines and proteases, such as IL-1, IL-6, VEGF, IL-8, CXCL9, and MMPs, are released, is one of the most characterized phenomena in senescence. The transcription factor GATA4 is degraded with the association of the autophagic adaptor p62 by selective autophagy under normal conditions, whereas the DNA damage response (DDR) kinases ataxia telangiectasia mutated (ATM) and ataxia telangiectasia and Rad3-related (ATR) receive senescence signals to facilitate the dissociation between GATA4 and p62 and stabilize GATA4, in turn activating NF-kB through tumor necrosis factor receptor-associated factor interacting protein 2 (TRAF3IP2) and IL1A(Kang *et al*, 2015). SASP is completely impaired in ρ0 cells by another method that does not use ethidium bromide but instead forces mitophagy(Correia-Melo *et al*., 2016). Mitochondrial disease patient-derived fibroblasts showed SASP at an early PDL and a short lifespan, whereas MirCs derived from the diseased cells partially depressed SASP and significantly extended their lifespan, suggesting that the antagonistic aging hallmarks might depend upon mitochondrial DNA.

It remains to be solved how replication of the mammalian mitochondrial genome is regulated for homeostasis and how heteroplasmy is propagated. MRT presented a concern, “reversion”, which occurs in embryonic stem cells derived from the embryo receiving the technology(Greenfield *et al*., 2017). A significant number of these ES cell lines showed the gradual domination of the karyoplast-associated mtDNA with time in culture to reach homoplasmy, called reversion(Hyslop *et al*, 2016; Kang *et al*, 2019; Yamada *et al*., 2016), although oocyte karyoplasts carried over less than 1% in maternal spindle transfer of Rhesus macaque(Lee *et al*, 2012). *C. elegans* demonstrated that the mitochondrial unfolded protein response (UPRmt) functions to maintain a heteroplasmy and propagates mutated mtDNA following a disturbance of the heteroplasmy to recover the original heteroplasmy(Lin *et al*, 2016). Prior to the study, UPRmt was only considered to promote the functional recovery of damaged mitochondria(Nargund *et al*, 2015); however, it is now understood as a double-edged sword, relying upon the quality of mtDNA (Tian *et al*, 2016). The preservation of mutated mtDNA might be a selfish aspect to survive. Although mammalian cells possess the same molecular machinery is currently unknown, the entity of the compatible machinery in mammals might be highly probable(Fiorese *et al*, 2016).

In addition to mitochondrial gene editing using ZFN(Gammage *et al*., 2016) and TALEN(Bacman *et al*., 2013), a sophisticated study was recently presented that designed a split new type of cytidine deaminase (DddA) combined with TALE protein to recognize a defined DNA sequence in tandem of mitochondrial targeting sequence (MTS) and separately uracil glycosylase inhibitor (UGI) to prevent the conversion of uracil to cytidine, called DdCBE(Mok *et al*, 2020). One concern is that DdDBE reduced the copy number of mtDNA, as MirC generation where it takes a period to reconstitute a new mitochondrial complex and network (Fig 3B). When this technology is applied in vivo, the cells that received the gene transfer could not be permissive due to a serious shortage of energy. In this DdCBE, uracil glycosylase inhibitor (UGI) functions to protect uracil to revert back to cytidine, leading to introduction of adenine in the counter strands upon the next round of mtDNA replication, therefore theoretically changing the heteroplasmy to 50% at maximum. In addition, DdCBE could not target deletions and long-range replacement. MirC provides advantages of superior heteroplasmy changes to dominate over the previous genotype and applicability to any mutation sequences, including deletions and long-range replacement, as well as no concern regarding off-target risks.

This somatic cell protocol offers drastic metabolic and lifespan changes through the combination of a decrease in mtDNA via a mitochondria-targeted endonuclease and the coincubation of isolated mitochondria. The replacement of mtDNA, which is regulated by nutrient sensors, is stable without reversion and is achieved by horizontal gene transfer in the cytoplasm of eukaryotes. MirCs could be clinically applied to treat not only mitochondrial diseases but also neurodegenerative diseases and cancers.

## Materials and Methods

### Plasmid production and Transfection

The mitochondrial targeting sequence fused with the 5’-end of DsRed2 (MTS-DsRed2) was digested from the pDsRed2-Mito vector (Clontech Laboratories, Inc., Palo Alto, CA, USA) with restriction enzymes and inserted into the pMXs-puro retroviral vector (Cell Biolabs, Inc. San Diego, CA. USA). The pCAGGS-MTS-*Xba*IR-P2A-PuroR and pCAGGS-MTS-EGFP-P2A-PuroR plasmids were synthesized from Genewiz (Suzhou, China). The DNA sequence of *Xba*IR referred to AF051092 of NCBI GenBank. These plasmids were transfected into NHDFs and 7SPs by electroporation using the Amaxa nucleofector kit and nucleofector 2b (Lonza, Walkersvill, MD, USA). Two days later, MTS-*Xba*IR- or MTS-EGFP-transfected cells were purified with 2 μg/ml puromycin treatment for 24 h. The plasmids used to assess the heteroplasmy ratio of mitochondria synthesized the sequence corresponding to the TaqMan probes and inserted them directly by GeneWiz (South Plainfield, NJ, USA).

### Human fibroblast culture

NHDFs were obtained from Lonza (Walkersvill, MD, USA). Leigh syndrome patient-derived skin (7S) fibroblasts were kindly provided by Koinobori Associate Inc., which supports research on mitochondrial diseases under approval from the ethical committees of our institution and Koinobori Associate Inc. Human uterine EMCs (EPC100) were kindly provided by Dr Umezawa (National Center for Child Health and Development). NHDFs and EPC100 were maintained in high-glucose DMEM (043-30085, Fujifilm Wako Pure Chemical, Osaka, Japan) supplemented with 10% fetal bovine serum and 1% penicillin/streptomycin (Thermo Fisher Scientific incorporated, Waltham, MA USA). The 7SPs were maintained in low-glucose DMEM with pyruvate (11885084) supplemented with 10% fetal bovine serum (FBS) and 1% penicillin/streptomycin (Thermo Fisher Scientific incorporated). All cell lines were incubated at 37°C under 5% CO_2_ and cultured at ∼80% confluence.

### Cell counting and cell diameter measurement

Cell number and cell viability were measured using the automatic cell counter ADAM (NanoEnTek Inc., Seoul, Korea), according to the manufacturer’s recommendations. Cell diameters were measured using the Scepter 2.0 Cell Counter and Scepter Software Pro (Merck KGaA, Burlington, MA, USA), according to the manufacturer’s recommendations.

### Mitochondrial isolation and transfer to human fibroblasts

Mitochondria were isolated from MTS-DsRed2 EMCs by differential centrifugation as described previously. In brief, the cells were harvested from culture dishes with homogenization buffer [HB; 20 mM HEPES-KOH (pH 7.4), 220 mM mannitol and 70 mM sucrose] containing a protease inhibitor mixture (Sigma-Aldrich, St. Louis, Missouri, USA). The cell pellet was resuspended in HB and incubated on ice for 5 min. The cells were ruptured by 10 strokes of a 27-gauge needle on ice. The homogenate was centrifuged (400 ×*g*, 4°C; 5 min) two times to remove the unbroken cells. The mitochondria were harvested by centrifugation (6000 ×*g*, 4°C; 5 min) and resuspended in HB. The amounts of isolated mitochondria were expressed as the protein concentration using a Bio-Rad protein assay kit (Bio-Rad Laboratories, incorporated, Richmond, CA, USA). Mitochondrial transfer was conducted by coincubating the isolated mitochondria with the plasmids or mRNAs transferred cells in 2 ml of standard medium at 37°C under 5% CO_2_ for 24 h.

### MtDNA digestion and quantitative assay

Total DNA was extracted from cells using NucleoSpin Tissue (Macherey-Nagel, Duren, Germany). The extracted DNA (100 ng) was digested with restriction enzymes (*Xba*IR or *Not*I) at 37°C for 30 min and subjected to selective amplification by PCR using KOD FX Neo (Toyobo Co., Ltd., Osaka, Japan) under the following conditions: 35 cycles (98°C for 10 sec, 60 °C for 30 sec and 68 °C for 30 sec) after initial denaturation (94 °C for 2 min). The primers used in this experiment are listed in Appendix Table S1. Reaction specificity was verified by agarose gel electrophoresis, and DNA bands were visualized using the ChemiDoc XRS+ System (Bio-Rad Laboratories, Incorporated).

MtDNA CN was analyzed by quantitative PCR. Quantitative PCR was performed using the extracted DNA (100 ng) as template with Kapa SYBR Fast qPCR Kit Master Mix (2×) Universal (Kapa Biosystems Ltd., Wilmington, MA, USA) on a CFX connect real-time system (Bio-Rad Laboratories, Incorporated, Hercules, CA, USA) under the following conditions: 40 cycles of PCR (95°C for 10 sec, 60°C for 1 min and 72°C for 30 sec) after initial denaturation (95°C for 2 min). The primers used in this experiment are listed in Appendix Table S1.

### MtDNA mutation Heteroplasmy analysis

To determine mutation ratios, we designed wild-type and mutant allele-specific TaqMan probes for the TaqMan SNP assay. The extracted DNA (1 ng) was used for quantitative PCR with the TaqMan Universal PCR Master Mix kit (Thermo Fisher Scientific Incorporated) on a CFX connect real-time system (Bio-Rad Laboratories, Incorporated) under the following conditions: 40 cycles of PCR (95°C for 15 sec and 60°C for 1 min) after initial denaturation (95°C for 10 min). A calibration curve was created using known CNs of plasmids containing the amplified mtDNA ND3 fragments for either wild-type or mutant sequences (Appendix Table S1). The mtDNA CN was estimated from the content ratio of 12S rRNA on mtDNA and *ACTB* (or *Actb*) on nuclear DNA by delta cycle threshold-based relative quantification. The primers and probes used in this experiment are listed in Appendix Table S1.

### RNA isolation, reverse transcription PCR and quantitative PCR

Total RNA from cells was extracted using TRIzol (Thermo Fisher Scientific Incorporated) and a Direct-zol RNA MiniPrep Kit (Zymo Research, Irvine, CA, USA) with DNase I, according to the manufacturer’s recommendations. To perform the qRT-PCR assay, 100 ng of total RNA was reverse-transcribed using the PrimeScript RT Reagent Kit (Takara Bio, Shiga, Japan) and a T100 thermal cycler (Bio-Rad Laboratories, Incorporated). qRT-PCR was performed with Kapa SYBR Fast qPCR Kit Master Mix (2×) Universal (Kapa Biosystems Ltd., Wilmington, MA, USA) on a CFX connect real-time system (Bio-Rad Laboratories, incorporated). The primers used in this experiment are listed in Appendix Table S1. The relative gene expression levels of *EGFP* and *Xba*IR were normalized to human *GAPDH* or mouse *Gapdh* expression.

### Measurements of cellular Bioenergetics

An Oroboros Oxygraphy-2k (Oroboros Instruments, Innsbruck, Austria) was used to measure cellular bioenergetic changes in cells, as described previously. In brief, the cells were harvested in 1 × 10^6^ cells/2 ml culture media. The cell suspension was transferred to a well, and Oroboros oxygraphy-2k was used. After baseline measurements, oligomycin (2 μg/ml), carbonyl cyanide-p-trifluoromethoxyphenylhydrazone (FCCP, 1 μM), rotenone (0.5 μM) and antimycin A (2.5 μM) were sequentially added to each well. Data are expressed as the oxygen consumption rates (O_2_ flow per cell; pmol/sec/cell). Routine, basal respiration, ETS, free routine activity, ROX, proton leakage and routine coupling efficiency were calculated as described previously.

### Time-lapse fluorescent microscopy and high-resolution microscopy

To observe DsRed2-mitochondrial uptake into cells, time-lapse fluorescence microscopy was performed using the JuLI stage (NanoEnTek Inc.). Red fluorescent images and phase images were taken every 30 min for 144 h from the beginning of mitochondrial coincubation. Fluorescence intensity was analyzed using JuLI STAT software (NanoEnTek Inc.). High-resolution microscopy of mitochondria was performed using N-SIM S (Nikon Corporation, Tokyo, Japan).

### iPS cell generation

Five days after mitochondrial transfer, the cells were seeded on 6-well plates at 1 × 10^5^ cells per well. The next day, the four Yamanaka factors (OCT3/4, SOX2, KLF4, and c-MYC) were introduced into cells by the Sendai Reprogramming vectors. At day 6, the cells were harvested by trypsinization and plated onto feeder-MEF (ReproCELL, Kanagawa, Japan) at 6 × 10^5^ cells per 60 mm dish. After 24 h, the medium (DMEM containing 10% FBS) was replaced with iPSC culture medium for primate embryonic stem (ES) cell medium (ReproCELL) supplemented with 20 ng/ml basic fibroblast growth factor (bFGF: Fujifilm Wako Pure Chemical). Approximately two weeks later, some granulated colonies appeared. At day 23, ES cell-like colonies were picked with a 200 μl tip and transferred to a 6-well plate containing adhered feeder-MEF. These colonies were stained using the BCIP/NBT Substrate System (Agilent Technologies, Santa Clara, CA, USA) according to the manufacturer’s recommendations. Alkaline phosphatase (AP)-positive colonies were counted using ImageJ (National Institutes of Health, Bethesda, MD, USA).

The well-grown colony lines, such as ES cells, were maintained by changing the iPS culture medium daily and passaging the cells every 5–6 days. The cells were harvested with CTK solution [2.5% trypsin, 5 ml of 1 mg/ml collagenase-IV (Thermo Fisher Scientific incorporated), 500 μl of filtered (0.2 μm) 0.1 M CaCl_2_, and 10 ml of Knockout Serum Replacement (KSR: Thermo Fisher Scientific Incorporated)]. The harvested cells were transferred to 15 ml conical tubes and centrifuged at 160 ×g for 5 min. The supernatant was carefully discarded without disturbing the cell pellet. Then, the cells were resuspended with fresh iPS culture medium and mixed by a soft twice to three times pipetting. Slightly crushed colonies were plated onto a new feeder-MEF-coated 60 mm dish. The feeder-MEFs were seeded at 3 × 10^5^ cells per gelatin-coated 60 mm dish on the day before passage.

### Immunocytochemistry

The cells were fixed with 4% paraformaldehyde at 4°C for 5 min and permeabilized with 0.1% Triton X-100 at room temperature for 20 min in the presence of a protein-blocking solution consisting of PBS supplemented with 5% normal goat serum (Agilent Technologies, Inc., Santa Clara, CA, USA). The cells were incubated overnight with primary antibodies in PBS at 4°C. They were washed extensively in PBS and incubated at room temperature for 30 min with secondary antibody. The nuclei were counterstained with 4’,6-diamidino-2-phenylindole (DAPI; diluted 1:500; FUJIFILM Wako Pure Chemical) in PBS at room temperature for 30 min. To prevent fading during microscopy, the cells were mounted in DakoCytomation Fluorescent Mounting Medium (Agilent Technologies, Inc.). Immunofluorescence images were visualized and recorded using a Biorevo BZ-9000 fluorescence microscope (Keyence Corporation, Osaka, Japan).

### Evaluation of plasmid transfection efficiency

The transfection efficiency of the plasmid was quantified by calculating the expression rate of EGFP. The expression of EGFP was measured before and after treatment with puromycin treatment for 24 h. After 30 min of incubation in medium supplemented with Hoechst 33342 (Dojindo Molecular Technologies, Inc., Kumamoto, Japan), fluorescent images were captured at random 3 points using an IX71 fluorescence microscope (Olympus, Tokyo, Japan). Transfection efficiency was determined by the ratio of EGFP-positive cells to Hoechst 33342-positive cells.

### Immunoassay and flow cytometric analysis for AMPK and S6 in ρ^-^ NHDFs

Six days after plasmid transfection, we performed a protein immunoassay using antibodies against AMPK, pAMPK, S6, and pS6 on a Wes system (ProteinSimple, Inc., San Jose, CA, USA). We compared the activation levels of AMPK and S6 in ρ^-^ NHDFs with other conditions: rapamycin, phosphatidic acid and starvation. Briefly, cells were treated with 50 nM rapamycin (Merck Millipore, Billerica, MA, USA) or 20 μM phosphatidic acid (Avanti Polar Lipids, Alabaster, Alabama, USA) for 24 h prior to sample collection. Starvation was performed by incubation with amino acid-free DMEM (Fujifilm Wako Pure Chemical) for one hour before sample collection.

For the immunoassay, treated cells were washed with PBS and collected into RIPA buffer (Fujifilm Wako Pure Chemical) containing protein inhibitor (Sigma-Aldrich). The cells were incubated on ice for 30 min and then homogenized. The extracts were spun down at 10,000 ×g for 10 min at 4°C, and the supernatants were analyzed using a Bio-Rad protein assay kit (Bio-Rad Laboratories, incorporated). The protein concentration of all samples was 0.2 mg/ml, and the primary antibody was diluted 50 times. The primary antibodies used in this experiment are listed in Appendix Table S1.

For flow cytometric analysis, cells were washed with PBS, harvested with 0.25% trypsin-EDTA and subjected to FCM analysis. The DsRed2-positive cell population was evaluated using 488 and 561 nm lasers. Fluorescence data were collected using SH800S (Sony). The flow cytometry files were analyzed using FlowJo software (Becton, Dickinson and Company).

### Evaluation of tolerance to H_2_O_2_ stimulation and starvation

We compared tolerance to H_2_O_2_ stimulation and starvation between each cell using the Annexin V-FITC Apoptosis Detection Kit (Nacalai Tesque, Kyoto, Japan). Cells were seeded in 6-well plates at 1 × 10^5^ cells per well. The next day, 600 μM H_2_O_2_ (Fujifilm Wako Pure Chemical) or amino-acid-free DMEM (Fujifilm Wako Pure Chemical) was added to the cells. After 3 h H_2_O_2_ treatment or 48 h starvation, the cells were washed with PBS and collected in centrifuge tubes. Annexin V-FITC and PI solution were added to the cells and allowed to react for 30 min at room temperature protected from light. Then, the cells were rapidly analyzed by FCM analysis using SH800S (Sony) and FlowJo software (Becton, Dickinson and Company).

### STR profiling

A GenePrint 10 System PCR Amplification kit (Promega Corporation) was used to determine the genetic signature of both samples based on the multiplex analysis of 9 loci and the Amelogenin sex-determining marker. PCR products were run in the Applied Biosystems 3730xl DNA Analyzer (Thermo Fisher Scientific Incorporated) and analyzed using GeneMapper ID Software (Thermo Fisher Scientific Incorporated) following the manufacturer’s recommendations.

### Single-cell droplet digital PCR (sc-ddPCR)

The sc-ddPCR protocol commenced with the encapsulation of a single cell into one oil droplet and then proceeded to the PCR step with a set of primers and fluorescent probes, which were the same as those used in the TaqMan SNP genotyping assay mentioned above, using TaqMan Polymerase with a 5’ to 3’ exonuclease, which releases the fluorophore from the probe, followed by the detection of the fluorescent signal in the droplets. The PCR mixture consisted of 4 μl of resuspended cells at a concentration of 2.5×10^5^/ml or 1.25×10^5^/ml, 10 μl of 2×ddPCR Supermix (Bio-Rad), wild-type and mutant allele-specific TaqMan probes at a concentration of 0.25 μM, primer mixtures at a concentration of 0.9 μM for the target gene, and nuclease-free water for a final volume of 20 μl. Droplets were generated using the Bio-Rad QX200 system (Bio-Rad) following the manufacturer’s instructions. The reactions were transferred to a 96-well plate (Eppendorf Corp., Hamburg, Germany) for PCR using a thermal cycler (Bio-Rad) under the following conditions: amplification was carried out at a regular ramp rate of 2.0°C /sec at 95°C for 10 min followed by 40 cycles of 30 sec at 95°C plus 2 min at 56°C. The final enzyme deactivation step occurred at 98°C for 10 min. The 96-well plate was transferred to a QX200 Droplet Reader (Bio-Rad), and the number of fluorescent droplets was analyzed. Each droplet was analyzed individually using a two-color detection system (set to detect FAM and VIC). The fluorescent droplets were counted to provide an absolute quantification of target mtDNA in digital form using QuantaSoft software 1.7 (Bio-Rad).

### Statistical analysis

All calculation was performed and plots were created using Prism 8 (GraphPad Software Inc., San Diego, CA, USA). The results are expressed as the mean ± S.E. The statistical significance of differences between the groups was evaluated using Student’s t-test, and P-values < 0.05 were considered significant.

## Data availability

Raw data are available on request. Expanded View for this article is available online.

## Acknowledgements

We would like to express our sincere thanks to Ms. Sayuri Shikata for her technical assistance and Mr. Taro Inaba, who is a managing director of REMIGES Ventures, for meaningful discussion regarding this study. Osaka, who is a professor in the Department of Pediatrics, Jichi Medical University, Japan, for providing patient-derived cells and the patient who donated her tissues to him.

## Funding

Funding was provided by a collaborative research grant from IMEL Biotherapeutics, Inc.

## Author contributions

S.G. designed the research; H.M. and D.K. performed all experiments; R.M. and A.S. analyzed and discussed the data; and S.G., H.M., D.K. wrote the paper.

## Conflict of interest

S.G. and D.K. received a collaborative research grant from IMEL Biotherapeutics, Inc. H.M., R.M. and A.S. have no competing interests.

## Supporting Information

Appendix

Movie EV1-1 to EV1-4

Movie EV2-1 to EV2-4

Movie EV3-1 to EV3-4

Movie EV4

